# Cross-Kingdom Control: Yeast Prion Protein [*MRPL10*^+^] Modulates Host Physiology in *Drosophila*

**DOI:** 10.64898/2026.08.26.747210

**Authors:** Chih-Chun Janet Lin, Jessica Y. Jiang, Maushmi D. Chitale, Elissa J. Cosgrove, Alexandria Van Elgort, Asha M. Jain, Julia C. Kelso, Xinyue Cui, Nilay Yapici, Andrew G. Clark

## Abstract

Prions, once mainly studied for their pathogenic roles, are now gaining recognition as adaptive elements in microbial physiology. Over one-third of wild yeast isolates harbor prion proteins, yet their impact on host-microbe interactions remains poorly characterized. Given the ecological dominance of yeasts in the *Drosophila* mycobiome, we leveraged the *Drosophila melanogaster*—*Saccharomyces cerevisiae* system to investigate how the mycobiome-derived prion, [*MRPL10^+^*], modulates host physiology. We show that flies exposed to [*MRPL10^+^*] yeast exhibit significantly enhanced cold tolerance and increased locomotor activity. This effect persists with heat-killed yeast and diluted culture, suggesting a stable, potent bioactive factor. Using the genetically diverse *Drosophila* Global Diversity Lines (GDL), we identified natural variation in responsiveness to [*MRPL10^+^*] yeast. Genome-wide association and functional RNAi screening revealed a gut-brain signaling axis involving genes critical for digestion, intercellular communication, transcription regulation, and neural transmission. Notably, serotonin and octopamine pathways were essential for [*MRPL10^+^*]-induced changes in cold tolerance and locomotion, implicating neuromodulatory circuits in prion-mediated microbial signaling. Our findings establish a mechanistic link between a fungal prion and host metabolic and neural adaptation. This work provides the first genetic dissection of a prion-mediated host-microbe interaction, laying the groundwork for investigating beneficial prions in complex microbial communities and highlighting a new dimension of the mycobiome’s influence on animal physiology.

## Introduction

Prion proteins are not always detrimental. In yeast, prions such as [*PSI*^+^], [*URE3*], and [*MOT3*^+^] take on self-propagating conformations that confer heritable phenotypes that are epigenetically transmitted, independent of DNA sequence changes. These conformational changes reshape cellular physiology, facilitating survival and environmental stress adaptation (Alberti et al., 2009; Chernova et al., 2014; Halfmann et al., 2010; Halfmann et al., 2012; Saad & Jarosz, 2021). Prion-driven traits contribute to gene regulation (Alberti et al., 2009; Du et al., 2008; Patel et al., 2009; Rogoza et al., 2010; Wickner, 1994), stress resistance (Suzuki et al., 2012; Tyedmers et al., 2008), and metabolic reprogramming (Du et al., 2008; Jarosz et al., 2014), and may even accelerate evolutionary processes (Halfmann et al., 2012; Lancaster et al., 2010; Lancaster & Masel, 2009).

Given these findings in yeast, it is plausible that prion-mediated changes could extend beyond the microbial cell. Here, we propose a novel hypothesis: a fungal prion can influence not only fungal physiology but also host biology and behavior, extending its effects from a cell-autonomous phenomenon to a cross-species signaling mechanism. The fungal microbiome—or mycobiome—is an integral, yet underexplored, component of the host ecosystem and is increasingly recognized for its roles in health and disease (Gaspar et al., 2025; Hill & Round, 2024; Huang et al., 2024). In clinical isolates of *Saccharomyces cerevisiae*, stress-induced aggregation of prion-like proteins such as Cdc19, Yef3, and Nop1 has been linked to drug resistance traits (Chen et al., 2021), potentially influencing mycobiome composition and thereby affecting host physiology. However, direct evidence connecting mycobiome-derived prions to host phenotypes remains scarce, and the underlying molecular mechanisms are poorly understood.

The *Drosophila* mycobiome naturally contains yeasts like *Saccharomyces* and *Candida* (Jones et al., 2022; Majumder et al., 2020; Quan & Eisen, 2018). Yeast serves not only as a nutritional source but also as a reservoir of signaling molecules that regulate host fly development (Grangeteau et al., 2018; Jimenez-Padilla et al., 2024), reproduction (Billeter & Wolfner, 2018), behavior (Christiaens et al., 2014; Murgier et al., 2019; Steck et al., 2018), metabolism (Colinet & Renault, 2014; Steck et al., 2018), thermal stress tolerance (Colinet & Renault, 2014), and lifespan (Libert et al., 2007). Notably, *Drosophila* also facilitates the spread of yeast by transporting fungal species across environments and aiding their colonization (Buser et al., 2014; Cho & Rohlfs, 2023; Christiaens et al., 2014; Korkmaz et al., 2025). This intimate and reciprocal ecological relationship makes the *Drosophila–Saccharomyces* system an ideal system for studying fungal-host interactions. Importantly, the availability of robust genetic toolkits for both yeast and fruit flies, along with the ability to rear flies under axenic or gnotobiotic conditions (Koyle et al., 2016), makes this system uniquely tractable for mechanistically dissecting how fungal prion states may influence host physiology and behavior.

## Results and discussion

### Remarkable cold tolerance induced by [*MRPL10^+^*] prion yeast in flies

To identify yeast-derived prions that influence host physiology, we screened a panel of prion-positive yeast strains for their effects on *Drosophila*. These yeast strains were isolated from a transient overexpression screen for stable, heritable prion traits (Chakrabortee et al., 2016). We found the [*MRPL10^+^*] prion strain conferred a pronounced cold tolerance phenotype (Supplementary Figure 1A). Flies exposed to [*MRPL10^+^*] yeast exhibited > 8-fold greater resistance to chill coma after a 2-hour cold shock compared to germ-free or prion-free yeast controls (Supplementary Figure 1A and Figure 1A). This enhancement was specific to acute cold shock, with no effect on long-term cold survival (Supplementary Figure 1B and Figure 1B). In contrast, diverse wild isolates from the Saccharomyces Genome Resequencing Project (SGRP) collection showed no comparable cold tolerance improvement (Supplementary Figure 1C). These findings demonstrate that the enhanced host cold tolerance is a specific property of the [*MRPL10^+^*] strain, not a general feature of prion-containing or wild yeast.

**Figure 1.**
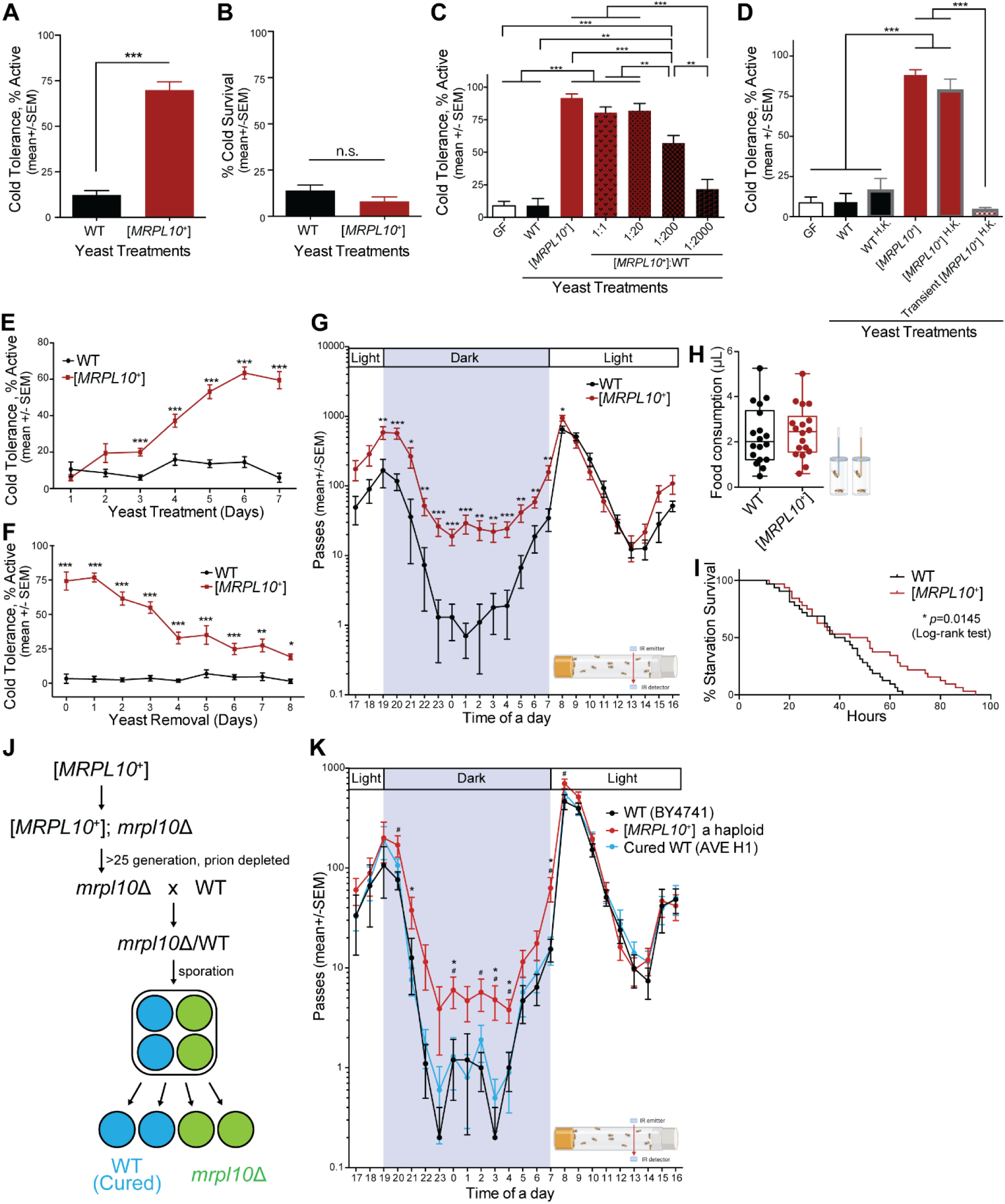
[*MRPL10*^+^] prion yeast modulates cold tolerance and locomotion in *D. melanogaster* via stable and reversible mechanisms. (A) Flies fed with [*MRPL10*^+^] yeast exhibit significantly enhanced cold tolerance following a 2-hour exposure to 4–6 °C, compared to those fed with wild-type yeast (BY4743; used as the default wild-type unless otherwise noted). (B) No significant differences in survival were observed between [*MRPL10*^+^]- and wild-type– treated flies after 3 days at 4–6 °C. (C) The cold-tolerance–promoting activity of [*MRPL10*^+^] yeast persists even at a 1:200 dilution, significantly exceeding that of wild-type and germ-free (GF) controls. (D) Heat-killed (H.K.) [*MRPL10*^+^] yeast also promote cold tolerance. This effect is reversible: flies lose the cold-tolerant phenotype after 6 days of exposure to heat-killed [*MRPL10*^+^] yeast followed by 4 days on GF conditions. (E) A time-course analysis reveals that [*MRPL10*^+^] yeast requires at least 3 days to induce cold tolerance, with effects plateauing after 6 days. (F) The cold-tolerance effect remains detectable, though reduced, even after removal of live [*MRPL10*^+^] yeast. (G) [*MRPL10*^+^]-fed flies display significantly elevated locomotor activity, particularly during the night phase. (H) No significant differences in food intake were observed between flies fed wild-type or [*MRPL10*^+^] yeast. (I) Flies fed with [*MRPL10*^+^] yeast exhibit greater resistance to starvation than those fed wild-type yeast. (J) Schematic of the prion-curing process used to eliminate [*MRPL10*^+^] from yeast, generating the cured strain. (K) The nighttime hyperactivity phenotype induced by [*MRPL10*^+^] yeast is reversed upon feeding with the cured yeast strain. All measurements in (A)-(I) and (K) are from three independent biological replicates.

The bioactive signal from [*MRPL10^+^*] yeast demonstrates exceptional potency and stability. Serial dilution assays revealed that even a 1:200 ratio of [*MRPL10^+^*] to prion-free isogenic yeast significantly enhanced cold tolerance in flies (Figure 1C), indicating a specific signaling mechanism rather than a general nutritional contribution. Notably, exposure to volatile compounds alone failed to induce cold tolerance (Supplementary Figure 2A), indicating that direct ingestion is required. Heat-killed [*MRPL10^+^*] yeast retained full bioactivity (Figure 1D), demonstrating heat stability of the active component. However, transient exposure to heat-killed [*MRPL10^+^*] yeast followed by transfer to germ-free conditions abolished the cold tolerance phenotype (Figure 1D), demonstrating that continuous exposure to [*MRPL10^+^*] yeast or its components is necessary to maintain the effect.

Time-course experiments revealed progressive development of [*MRPL10^+^*]-induced cold tolerance. Flies began to show a significant increase in chill coma resistance after three days of exposure and reached a plateau by day six (Figure 1E). Upon removal from live [*MRPL10^+^*] yeast, cold tolerance declined gradually yet remained elevated relative to non-prion controls (Figure 1F). The reversibility of heat-killed yeast effects versus the persistence with live yeast suggests that live yeast may trigger sustained changes beyond immediate dietary effects, possibly involving gut-associated fungal dynamics.

Chill coma—a temporary neuromuscular shutdown during cold stress (Macmillan & Sinclair, 2011)—prompted investigation into whether [*MRPL10^+^*]-induced cold tolerance enhances locomotor resilience. Surprisingly, [*MRPL10^+^*]-exposed flies exhibited significantly increased locomotor activity, with pronounced nocturnal enhancement (Figure 1G), a pattern typically associated with foraging and starvation responses. However, these flies were not starved; they maintained normal food intake (Figure 1H), showed no dietary avoidance for [*MRPL10^+^*] yeast (Supplementary Figure 2B), displayed heightened starvation resistance (Figure 1I), and retained full reproductive capacity (Supplementary Figure 2C). This rules out compensatory starvation response and instead suggests that [*MRPL10^+^*] promotes a state of enhanced neuromuscular readiness. To establish causality, we cured the [*MRPL10^+^*] yeast strain by eliminating the prion, preserving the genomic background (Figure 1J). Flies fed the cured yeast returned to baseline levels observed in prion-free controls (Figure 1K), confirming that the [*MRPL10^+^*] prion itself drives this novel host behavioral adaptation.

### Natural genetic variation reveals a gut-brain axis mediating microbial control of cold tolerance

The *Drosophila* Global Diversity Lines (GDL)—a genetically diverse panel derived from wild populations across five continents—capture extensive natural genetic variation (Grenier et al., 2015). These lines display normal development and viability, with no apparent feeding defect. To determine how host genetic background influences responsiveness to prion-derived microbial signals, we exposed GDL flies to [*MRPL10^+^*] yeast. We observed pronounced response heterogeneity: some lines exhibited a strong cold tolerance phenotype in response to [*MRPL10^+^*] and were classified as Type I responders (Figure 2A). This indicates that the effect is not restricted to a single laboratory strain but may reflect a more generalizable host–microbe interaction. In contrast, other lines showed little to no change in cold tolerance—either maintaining consistently high (Type II) or low (Type III) cold tolerance regardless of yeast condition (Figure 2A). These findings reveal that the [*MRPL10^+^*]-induced cold tolerance is not a universal trait but is shaped by host genetic background. Response magnitudes were quantitatively measured and ranked (Figure 2B), establishing a phenotype continuum for genome-wide association studies (GWAS) to identify host genes or pathways that govern sensitivity to prion-derived microbial signals and uncover key regulators of microbe–host communication.

**Figure 2.**
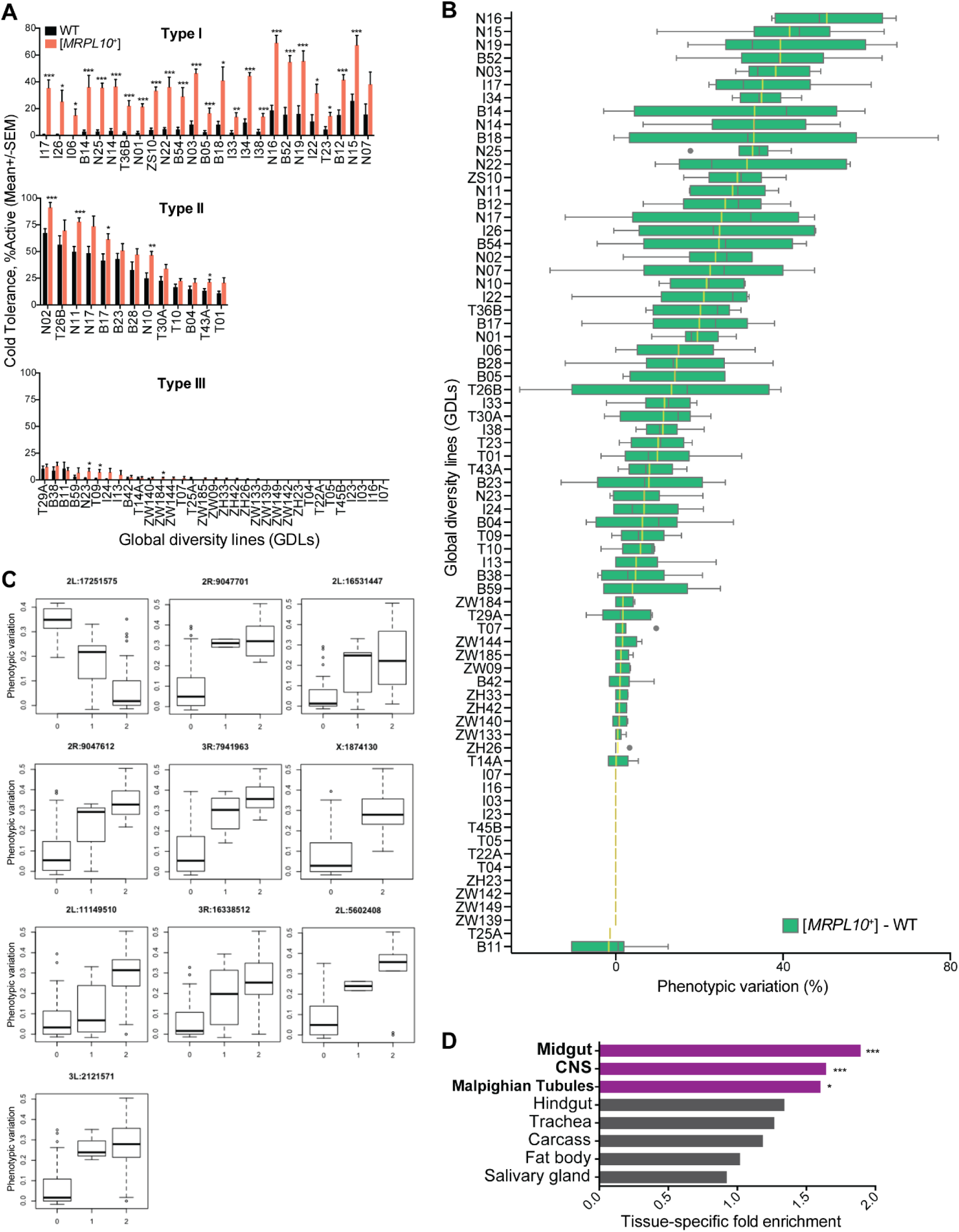
Natural variation in cold tolerance and genome-wide association mapping identify key loci and tissues mediating [*MRPL10*^+^]-dependent adaptation. (A) *Drosophila* Global Diversity Lines (GDL) exhibit distinct cold tolerance profiles in response to [*MRPL10*^+^] yeast, classified into three phenotypic response types: (I) differential response between [*MRPL10*^+^] and wild-type yeast, (II) non-responsive with constitutively high cold tolerance, and (III) non-responsive with constitutively low cold tolerance. (B) GDL strains ranked by phenotypic variation (difference in mean cold tolerance between yeast treatments), used as a quantitative trait for genome-wide association studies (GWAS). Tukey boxplots depict the median, interquartile range, and outliers (gray dots). The yellow line indicates the overall mean. (C) Top 10 single-nucleotide polymorphism (SNP) loci significantly associated with the cold tolerance response to [*MRPL10*^+^] yeast. Genotypes are represented on the x-axis as 0 (homozygous reference, 0/0), 1 (heterozygous, 0/1), and 2 (homozygous alternate, 1/1). The y-axis shows the corresponding phenotypic variation. Genomic coordinates are based on the *Drosophila melanogaster* reference genome (release 5.34). (D) Candidate genes linked to [*MRPL10*^+^]-induced cold tolerance are enriched in the midgut, central nervous system (CNS), and Malpighian tubules. All data in (A)–(C) are based on at least three independent biological replicates.

Genome-wide association analysis identified 145 variants (144 SNPs and 1 indel) significantly associated with [*MRPL10^+^*]-induced cold tolerance (BH-adjusted *p* < 0.1; Supplementary Table 1). These variants mapped to 246 candidate genes. The distribution and strength of associations are shown in the Manhattan and Q–Q plots (Supplementary Figure 3A and 3B). Notably, the top 10 SNPs exhibited substantial effect sizes on cold tolerance (Figure 2C). Functional enrichment analysis revealed significant over-representation of these genes in metabolic and neuronal pathways (Supplementary Figure 3C). Strikingly, these genes showed enriched expression in the midgut and central nervous system (Figure 2D), implicating a gut–brain axis in [*MRPL10^+^*]’s physiological effects.

**Figure 3.**
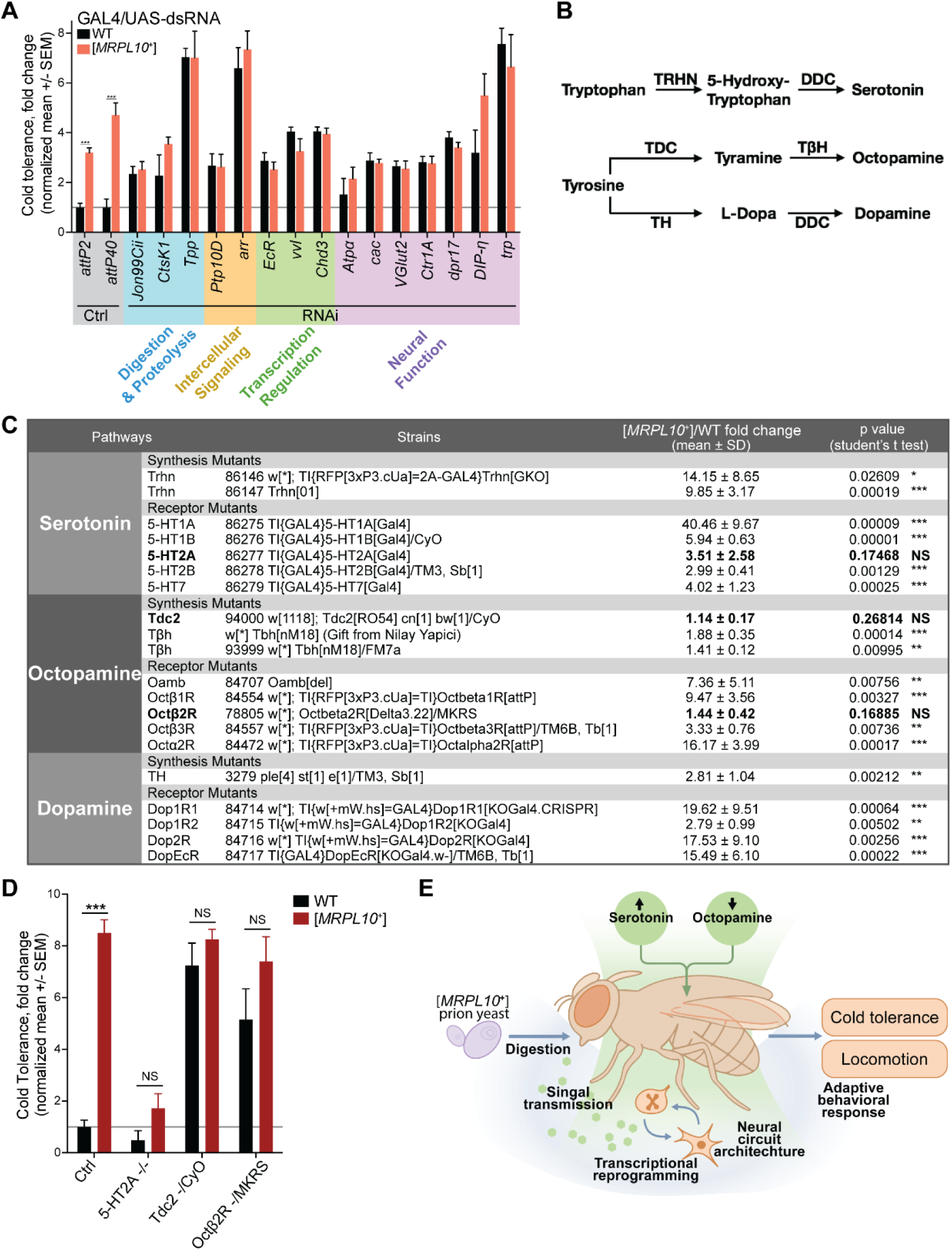
Functional genetic screen reveals gut-brain signaling and neuromodulation underlying [*MRPL10*^+^]-induced cold tolerance in *Drosophila*. (A) RNAi knockdown screening identified genes in four major functional categories required for the cold tolerance response to [*MRPL10*^+^] versus wild-type yeast: (1) digestion and proteolysis (*Jon99Cii*, *CtsK1* [also known as *26-29-p*], *Tpp* [*CG5171*]); (2) intercellular signaling and communication (*Ptp10D*, *arr*); (3) transcriptional regulation and chromatin remodeling (*EcR*, *vvl*, *Chd3*); and (4) neural function and neurotransmission (*Atpα*, *cac*, *VGlut2* [*MFS9*], *Ctr1A*, *dpr17*, *DIP-η*, *trp*). (B) Biosynthesis pathway of serotonin, octopamine, and dopamine in *Drosophila*. (C) Summary of *Drosophila* mutant lines screened for neuromodulatory components essential to the [*MRPL10*^+^]-induced cold tolerance phenotype. Fold changes and statistical significance (*p*-values) are reported for each genotype. (D) The differential cold tolerance responses to [*MRPL10*^+^] and wild-type strains are abolished in mutants lacking *5-HT2A*, *Tdc2*, or *Octβ2R*, indicating essential roles for serotonin and octopamine signaling. (E) Working model: [*MRPL10*^+^] yeast initiates a gut-brain signaling cascade involving digestive processes, signal transmission, and transcriptional reprogramming. This cascade converges on neural circuits requiring ion transport and synaptic function to mediate adaptive responses. Serotonin and octopamine pathways integrate these signals to coordinate host locomotion and cold tolerance.All data in (A), (C), and (D) are based on at least three independent biological replicates.

To functionally determine which of these GWAS candidate genes are required for the [MRPL10+]-induced phenotype, we performed an RNAi screen of 67 genes, selected based on GWAS significance, pathway enrichment, and availability of RNAi strains, using ubiquitous GAL4 drivers (Supplementary Table 2). Knockdown of 15 genes abolished [*MRPL10^+^*]-induced cold tolerance enhancement (Figure 3A, Supplementary Table 2), confirming their roles in mediating the host response. These genes cluster into functional modules:

- Digestive processors: *Jon99Cii* (serine protease), *CtsK1* (cysteine protease), and *Tpp* (Trehalose-6-phosphate phosphatase) suggest efficient nutrient metabolism enables prion signal detection.
- Signal transducers: *Ptp10D* (EGFR/FGFR/Pvr-regulating receptor tyrosine phosphatase) and *arr* (a transmembrane protein enabling Wnt receptor activity) implicate intercellular communication in microbial cue translation.
- Transcriptional regulators: *EcR* (ecdysone nuclear receptor transcription factor), *vvl* (POU domain homeobox transcription factor), and *Chd3* (nuclear ATP-dependent nucleosome remodeler) indicate hormonal/immune signaling and epigenetic reprogramming.
- Neural effectors: *Atpα* (Na+/K+ antiporter), *cac* (voltage-gated Ca^2+^ channel), *VGlut2* (glutamate transporter), *Ctr1A* (neuropeptide-maturing copper transporter), *trp* (a cation channel with high Ca^2+^-permeability), and synaptic complex *dpr17*/*DIP-η* involved in neuronal activity and synaptic function.

This multi-system involvement—spanning digestion, signaling, transcriptional regulation, and neural function—demonstrates how [*MRPL10^+^*] orchestrates host cold resilience through a gut-brain axis.

### Neuromodulatory control of cold tolerance and locomotion via serotonin–octopamine circuit dynamics

Given the functional associations of multiple neuronal genes and synaptic components, we next asked whether key neuromodulatory systems—serotonin, octopamine, and dopamine signalings—play a role in mediating the [*MRPL10^+^*]-induced cold tolerance. These biogenic amines are synthesized from amino acid precursors through well-characterized and evolutionarily conserved enzymatic pathways (Figure 3B): serotonin from tryptophan via *Trhn* and *Ddc*; octopamine from tyrosine via *Tdc2* and *Tbh*; and dopamine from tyrosine via *Th* and *Ddc* (Ozcete et al., 2024; Rosikon et al., 2023). Each neuromodulator acts through specific receptors to influence neural circuit activity. To functionally interrogate these pathways, we tested flies carrying mutations in key biosynthetic enzymes and receptors (Figure 3C).

Our functional screen identified three critical regulators: serotonin receptor *5-HT2A*, octopamine biosynthetic enzyme *Tdc2*, and octopamine receptor *Octβ2R*. Genetic disruption of *5-HT2A* completely abolished the [*MRPL10^+^*]-induced cold tolerance (Figure 3D), establishing serotonin signaling as essential for the phenotype. Conversely, loss of *Tdc2* or *Octβ2R* resulted in constitutively elevated cold tolerance even in the absence of [*MRPL10^+^*] exposure (Figure 3D), revealing that octopamine signaling acts as a negative regulator in this context. Together, these findings support a model in which [*MRPL10^+^*] yeast promotes cold tolerance by activating serotonin signaling while simultaneously suppressing octopaminergic signaling.

Since chill coma resistance reflects enhanced neuromuscular activity and resilience (Macmillan & Sinclair, 2011), we next dissect the neuromodulatory control of locomotion. Specific neuromodulatory neurons were silenced using GAL4 drivers crossed with UAS-TNT (White & Peabody, 2009). Silencing *ddc*+ (serotonergic and dopaminergic) and *tdc2*+ (octopaminergic) neurons abolished the differential locomotor activity normally observed between wild-type and [*MRPL10^+^*]-exposed flies (Supplementary Figure 4A). In contrast, silencing *th+* (dopaminergic-specific) neurons had no effect (Supplementary Figure 4A). These results demonstrate that serotonergic and octopaminergic circuits are key mediators of both [MRPL10^+^]-induced cold tolerance and locomotion enhancement.

Genetic perturbations further confirmed these pathways: mutations in the serotonin synthesis enzyme *Trhn* or serotonin receptors *5-HT1A*, *5-HT1B*, *5-HT2A*, and *5-HT2B* significantly reduced the [*MRPL10^+^*]-induced locomotor enhancement (Supplementary Figure 4B). Conversely, mutations in the octopamine synthesis enzyme *Tdc2* or octopamine receptors *Octβ2R* and *Octβ3R* constitutively elevated locomotion even in the absence of [*MRPL10^+^*] (Supplementary Figure 4C), suggesting that octopaminergic signaling normally restrains this behavior. In contrast, dopamine signaling proved to be largely dispensable; neither the biosynthesis enzyme *Th* nor most dopamine receptors had an effect, with the exception of Dop2R (Supplementary Figure 4D), suggesting potential pathway cross-talk or baseline modulation.

Collectively, these findings demonstrate that [*MRPL10^+^*] yeast enhances locomotion through dual neuromodulatory control: serotonin pathway activation coupled with octopaminergic suppression. The distinct receptor requirements for locomotion versus cold tolerance point to circuit-level specificity. We propose that cold tolerance is mediated by targeted serotonin–*5-HT2A* and octopamine–*Octβ2R* pathways, likely modulating central or peripheral systems governing chill coma dynamics. In contrast, locomotor activity represents a more complex and distributed behavioral output, relying on broad serotonin/octopamine signaling across multiple neural circuits.

In summary, we establish that the [*MRPL10^+^*] prion yeast specifically reprograms *Drosophila* physiology through a previously unrecognized gut-brain signaling axis, enhancing cold tolerance and locomotor behavior. Host genetic variation revealed variable responsiveness, enabling GWAS identification of a core set of genes—spanning digestive enzymes, transcriptional regulators, and neural signaling components—that transduce microbial signals to engage neuronal circuits and orchestrate systemic physiological changes in the host. Functionally, we show that serotonin and octopamine act antagonistically: serotonin signaling is required for cold tolerance and locomotor enhancement, while octopaminergic pathways tonically suppress these behaviors (Figure 3E). This work establishes mycobiome-derived prions as microbial effectors capable of perturbing host neuromodulatory networks, providing a mechanistic framework for how microbial prion states shape complex host phenotypes via the gut-brain axis.

Gene-environment interactions (GxE) are key drivers of evolutionary adaptation to local microbial environments (Lazzaro et al., 2008; Mullinax et al., 2025; Unckless et al., 2016). The differential responsiveness of geographically distinct *Drosophila* populations to [*MRPL10^+^*] yeast suggests a co-evolved relationship between host genotypes and fungal signaling traits, opening an exciting research direction at the intersection of microbial memory and animal physiology. Given that fungi have evolved sophisticated strategies for dispersal—often leveraging animal hosts (Chaudhary et al., 2022; Christiaens et al., 2014; Korkmaz et al., 2025) —it is intriguing to speculate that prion-like states in commensal yeasts may enhance host neuromuscular readiness not just for its survival, but for mutual benefit. In this scenario, [*MRPL10^+^*] yeast could promote cold escape behaviors in flies, aiding both partners: flies increase their chances of finding food and warmth, while yeasts gain opportunities for dispersal across the landscape. This potential symbiotic adaptation highlights a compelling new layer of microbe–host communication with ecological and evolutionary implications.

## Materials and methods

### Drosophila strains and husbandry

Global Diversity Lines (GDL) had been maintained in the Clark lab since their isolation (PMID: 25673134). Fly strains used for RNAi screening were obtained from the Bloomington *Drosophila* Stock Center (BDSC) (see Supplementary Table 2). Mutant strains were sourced either from BDSC or kindly provided by Nilay Yapici (Figure 3C). Genetic tools for neuronal silencing via the tetanus toxin system were acquired from BDSC, including *DDC-GAL4* (BDSC #7010), *Tdc2-GAL4* (BDSC #9313), *TH-GAL4* (BDSC #8848), and *UAS-TNT* (BDSC #28837).

All flies were reared on glucose–yeast–agar medium (“D food”; recipe available at https://cornellfly.wordpress.com/d-food/) at 25 °C under a 12-hour light–dark cycle. For experiments using the virginator stock (BDSC #8846), developing larvae were exposed to a 37 °C water bath for 1.5 hours to selectively eliminate males, resulting in the emergence of virgin females.

### Yeast strains and culture

*Saccharomyces cerevisiae* wild-type strains (BY4741, BY4742, and BY4743) were kindly provided by Jeffrey A. Pleiss (Cornell). Prion-positive strains, [*MRPL10^+^*]-cured strains (AVE H1), and SGRP lines were generously provided by Daniel F. Jarosz (Stanford). All yeast strains were cultured in 0.22-µm–filtered YPD medium (Millipore #1375) at 30 °C until reaching exponential growth phase. Cultures were then dispensed onto sterilized fly D food and incubated at 25 °C for an additional 3 days before exposure to adult flies.

For heat-killed preparations, yeast cultures were pelleted by centrifugation and incubated at 70 °C for 2 hours to ensure complete inactivation prior to plating on sterilized fly D food. Heat-killed yeast was replenished regularly to maintain consistent availability throughout the treatment period.

### Axenic fly preparation

Fly eggs were collected over a 24-hour period on grape juice agar plates composed of 10% yeast extract (Sigma #Y1625), 3% agar (Fisher #BP1423), 30% Welch’s grape juice concentrate, and 0.05% methyl paraben. Plates were placed in cylindrical egg-laying chambers, and eggs were gently harvested using a paintbrush and water into 50 mL Falcon tubes. Samples were centrifuged at 3,000 rpm for 1 minute, and the supernatant was carefully removed with a vacuum-connected, autoclaved glass pipette to avoid disturbing the egg pellet.

To surface-sterilize the eggs, 45 mL of freshly prepared 10% bleach solution was added, and the tube was gently shaken for 30 seconds. Eggs were pelleted again by centrifugation (3,000 rpm, 1 min), and the bleach was removed. This bleaching step was repeated once. Eggs were then washed four times with sterile water containing 0.06% Tween-20 to eliminate residual bleach.

Sterilized eggs were resuspended in sterile water and transferred into autoclaved fly D food vials using wide-bore pipette tips. Vials were incubated at 25 °C to allow axenic eggs to hatch and develop. Care was taken to avoid overflooding, as excess water can lead to poor hatching or larval survival. To confirm sterility, a small subset of processed eggs was inoculated into LB or YPD medium and incubated to monitor for microbial contamination.

Fly D food was prepared in polypropylene (PP) vials sealed with foam plugs and sterilized by autoclaving at 121 °C for 20 minutes (standard liquid cycle). Vials were cooled on a shaker at room temperature to ensure uniform consistency and stored at 4 °C until use.

### Cold tolerance assessment

Following the designated yeast treatment period, vials containing 25 ± 5 adult flies were inverted and placed in a cold room (4–6 °C) for 2 hours to induce chill coma. After cold exposure, vials were gently tapped, and flies were assessed based on their behavioral state using the inner surface of the white foam plug as a contrasting background. Flies lying on their backs and unresponsive to tapping were classified as being in chill coma, whereas those standing or moving were considered awake. Each vial was scored twice, and the mean of the two scores was used for analysis. A higher percentage of awake flies indicated greater cold tolerance. All assays were conducted at the same time of day under uniform lighting to minimize circadian influences.

### Cold survival test

Following the designated yeast treatment period, vials containing 25 ± 5 adult virginator flies were inverted and placed in a cold room (4–6 °C) for 72 hours. After cold exposure, vials were returned to 25 °C for a 24-hour recovery period before assessing survival. Flies that were standing upright and responsive to gentle tapping were scored as alive, while immobile or unresponsive flies were considered dead. All assays were performed at a consistent time of day under uniform lighting conditions to minimize circadian variability.

### Locomotor activity measurement

Locomotor activity was recorded using a horizontal Locomotor Activity Monitor (Trikinetics LAM25H). After the designated yeast treatment period, vials containing 25 ± 5 adult flies were positioned horizontally in the monitor, with infrared beam sensors aligned to bisect the center of each vial. Vials were secured with rubber bands to prevent sliding during recording. Beam breaks, reflecting fly locomotion, were logged every 10 minutes using Trikinetics DAMSystem software (v3.10.7). Monitors were housed in an incubator set to 25 °C with 50–70% relative humidity under a 12 h:12 h light–dark cycle. Flies were given a 2-hour acclimation period prior to data collection. Activity was recorded continuously for 3 consecutive days. Raw beam break counts were binned into 60-minute intervals and normalized to 25 flies per vial. All assays were conducted in parallel using at least three (ideally six) biologically independent vials per condition to minimize environmental variability.

### Food consumption measurement

Food intake was quantified using the Capillary Feeder (CAFE) assay. Adult virginator flies were subjected to yeast treatment for 5 days prior to transfer into 50 mL Falcon tubes (6 ± 2 flies per vial). Each tube had a needle-punctured cap fitted with a calibrated 5 µL glass capillary (VWR #53432-706) containing liquid food composed of 10% heat-killed, weight-normalized yeast and 10% glucose in distilled water. To maintain humidity and prevent desiccation, a 2% agar pad and a piece of thick filter paper soaked in distilled water were placed at the bottom of each tube. Identical tubes without flies served as evaporation controls. Tubes were maintained at 25 °C with 50–70% relative humidity under a 12 h:12 h light– dark cycle. Flies were allowed a 22-hour acclimation period before data collection. Food consumption was measured by recording the decrease in capillary liquid levels at 7–13 hour intervals, followed by capillary replacement, over a 2-day period. Net intake was calculated by subtracting the average evaporative loss (from control tubes) from the total volume decrease in experimental tubes. Consumption values were normalized to 6 flies per 12-hour period per vial.

### Food preference assay

Food preference was assessed using a modified Capillary Feeder (CAFE) assay, similar to the food consumption measurement described above. Virginator flies naïve to prior yeast exposure were used. Each 50 mL Falcon tube (containing six adult flies aged 3–5 days) was fitted with two calibrated 5 µL glass capillaries (VWR #53432-706), each filled with a different yeast-conditioned liquid food composed of 10% heat-killed, weight-normalized yeast and 10% glucose in distilled water. A 2% agar pad and a piece of water-soaked filter paper were placed at the bottom of each tube to maintain humidity. Tubes were maintained at 25 °C with 50–70% relative humidity under a 12 h:12 h light–dark cycle. The positions of the two capillaries were alternated across replicates to eliminate positional bias. After a 22-hour acclimation period, liquid consumption from each capillary was recorded at 7–13 hour intervals and replenished with fresh capillaries over a 2-day period. Identical tubes without flies served as evaporation controls. Net consumption was calculated by subtracting average evaporative loss from the total volume decrease. Consumption values were normalized to a 12-hour period per vial, and food preference was expressed as the percentage of total intake from each food type.

### Olfactory Exposure Assay

To evaluate the effects of yeast-derived volatile compounds, a custom two-compartment olfactory chamber was constructed using standard *Drosophila* vials. The bottom compartment contained D food supplemented with one yeast treatment, into which virginator flies were introduced. A breathable foam plug, trimmed to approximately 0.5–1 cm in thickness, was inserted above the flies to restrict them to the lower compartment while allowing airflow. The upper compartment contained D food with a second yeast treatment, serving as a source of volatile cues without direct contact with the flies. The vial was sealed with a second foam plug to maintain containment and airflow. Phenotypic assessments were performed after 5 days of olfactory exposure.

### Reproductive assay

To evaluate the impact of yeast treatments on female fecundity, initial matings were conducted on standard D food for 2 days. After mating, males were removed, and mated females (10 per vial) were transferred to vials containing the designated yeast treatment. Flies were transferred to fresh yeast-treated vials daily, and reproductive output was quantified by counting the number of adult progeny that emerged from each vial.

### Starvation survival test

After 5 days of yeast treatment, individual virginator flies were transferred to vials containing a 2% agar pad and a piece of water-soaked filter paper to provide moisture without nutritional content. Each vial was positioned horizontally in a Locomotor Activity Monitor (Trikinetics LAM25H), with an infrared beam aligned to the center to detect movement. Vials were secured with rubber bands to prevent shifting during monitoring. The system was maintained at 25 °C with 50–70% relative humidity under a 12 h:12 h light–dark cycle. Locomotor activity was recorded at 1-hour intervals using DAMSystem software (v3.10.7) and monitored continuously until all flies ceased movement. The time of the final recorded beam break was used as a proxy for time of death. Survival curves were generated using Kaplan–Meier analysis, and statistical significance was assessed using log-rank tests.

### Genome-Wide Association (GWA) Analysis Using Global Diversity Lines (GDL)

To identify genetic variants associated with differential cold tolerance, we utilized the *Drosophila melanogaster* GDL as a reference panel. These lines capture a broad spectrum of natural genetic variation from geographically diverse populations. Axenic eggs from each line were reared on D food, and 1–3-day-old adults were subjected to yeast treatments (wild-type vs. [*MRPL10^+^*]) for seven days prior to cold tolerance testing. A total of 66 strains, with both phenotypic and genomic data available, were included in the GWA analysis.

Genotype data, including high-confidence SNPs and short indels, were obtained from publicly available whole-genome sequencing datasets aligned to the *Drosophila melanogaster* reference genome (release 5.34). Variants were filtered to remove invariant sites, sites with >10% missing calls, and sites with >2 genotype classes (i.e., multiallelic or complex sites), resulting in a final dataset of 4,820,530 SNPs and 414,904 indels for the analysis. Phenotypic variation—defined as the difference in mean cold tolerance between yeast treatments—was modeled using a linear mixed model implemented with the `relmatLmer()` function from the *lme4qtl* R package. Individual SNP genotype was included as a fixed effect, and strain identity as a random effect. To account for population structure, a genetic relatedness matrix (GRM) was specified via the `relmat` argument, calculated as the covariance of the mean-centered SNP genotype matrix.

The significance of genotype effects was assessed using Wald tests. Multiple testing correction was performed using the Benjamini–Hochberg procedure, identifying 144 significant SNPs and 1 indel with an adjusted *p*-value < 0.1. Manhattan and Q–Q plots were used to visualize the association signals. Coordinates of significant loci were converted from genome release 5 to release 6 (v6.28.99) using the FlyBase Sequence Coordinates Converter and annotated using SnpEff to infer potential functional relevance, resulting in 246 annotated candidate genes.

### Tissue-specific gene expression enrichment analysis

RNA-seq data from FlyAtlas2 were used to assess tissue-specific expression enrichment of GDL-GWAS candidate genes. This dataset includes normalized FPKM values for 17,247 genes across various *Drosophila* tissues. For each gene, a tissue-specific enrichment score was calculated as:

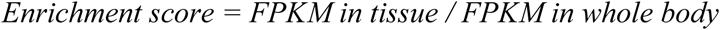

To avoid inflation from low denominator values, whole-body FPKM values <2 were set to 2, as recommended by FlyAtlas2. A gene was considered enriched in a given tissue if its enrichment score was ≥3, a threshold selected for statistical relevance.

Of the 246 GDL-GWAS candidate genes, 233 were annotated in the FlyAtlas2 dataset and thus included in the enrichment analysis. For each tissue, the fold enrichment of GDL-GWAS candidate genes relative to the background gene set was assessed using Fisher’s exact test in R (`fisher.test()` function), yielding odds ratios and associated *p*-values. Odds ratios were used as fold enrichment values for bar plot visualization.

|  | GDL-GWAS candidate genes | Background gene set |
| --- | --- | --- |
| Enrichment score $\geq 3$ | *a* | *N-a* |
| Enrichment score $< 3$ | *b* | *n-b* |
| Column total | *a+b=233* | *(N+n)-(a+b)=17,014* |

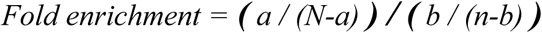

### Biological pathway enrichment analysis

To identify overrepresented biological processes among the 246 candidate genes from the GDL-GWAS, Gene Ontology (GO) enrichment analysis was performed using the Biological Process (BP) category. FlyBase gene IDs were submitted to the Database for Annotation, Visualization, and Integrated Discovery (DAVID), with *Drosophila melanogaster* specified as the background genome. The analysis was restricted to GO terms under the GOTERM_BP_DIRECT category. Enrichment significance was evaluated using Fisher’s exact test with default DAVID thresholds (Max.Prob. ≤ 0.1; Min.Count ≥ 2).

### RNAi-mediated knockdown in *Drosophila*

RNAi knockdown was carried out using the GAL4/UAS system. Males from either *Tubulin-GAL4* or *Armadillo-GAL4* ubiquitous driver lines were crossed to virgin females (4–10 days post-eclosion) carrying UAS-dsRNA constructs targeting genes of interest, at a ratio of 5 males to 15 females. Crosses were maintained at 25°C on standard fly D food and transferred to fresh vials every two days to ensure that F1 progeny were collected within a 2-day age window. F1 progeny expressing both the GAL4 driver and UAS-dsRNA transgene were subjected to yeast treatment (wild-type vs. [*MRPL10^+^*]) beginning at 2–4 days of adult age (25±5 flies per vial). Phenotypic assessments were conducted after 7 days of treatment, at 9–11 days of adult age.

### Neuron silencing using the tetanus toxin system

To functionally silence specific neuronal populations, the GAL4/UAS system was used to express the tetanus neurotoxin light chain (TNT), which blocks synaptic transmission by cleaving synaptobrevin. Male flies carrying *DDC-GAL4* (BDSC #7010), *Tdc2-GAL4* (BDSC #9313), or *TH-GAL4* (BDSC #8848)—which target serotonergic, tyraminergic/octopaminergic, and dopaminergic neurons, respectively—were crossed to virgin females carrying *UAS-TNT* (BDSC #28837). Progeny expressing both the GAL4 driver and *UAS-TNT* transgene served as the experimental group, while *UAS-TNT/Sb* siblings lacking GAL4 expression were used as negative controls. Flies were subjected to yeast treatments (wild-type vs. [*MRPL10^+^*]) beginning at 1–3 days of adult age and maintained on treatment for 6 days prior to phenotypic assessment.

---

**Supplementary Figure 1.**
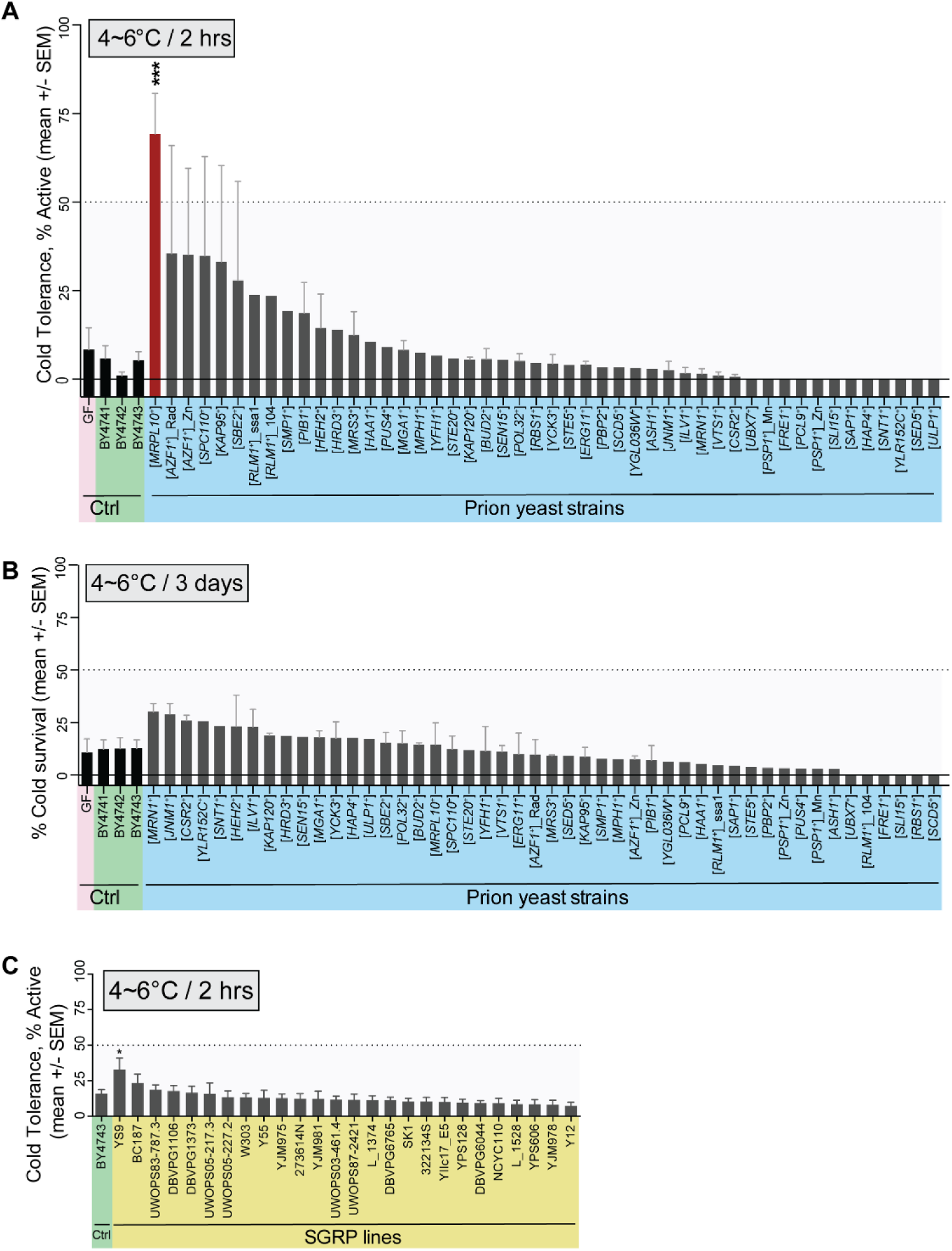
[*MRPL10*^+^] yeast confers superior cold tolerance in flies compared to other prion-positive and the SGRP yeast strains. (A) Flies fed with [*MRPL10*^+^] yeast exhibit significantly greater cold tolerance (4–6 °C for 2 hours) compared to germ-free control, wild-type yeast strains (BY4741, BY4742, BY4743), and other prion-positive yeast strains included in the screen. (B) No significant differences in cold survival (4–6 °C for 3 days) were observed across the various yeast treatments. (C) YS9, a yeast strain from the SGRP collection, modestly improved cold tolerance in flies, but its effect was notably weaker than that of [*MRPL10*^+^] yeast. All measurements in (A)-(C) are based on three independent biological replicates.

**Supplementary Figure 2.**
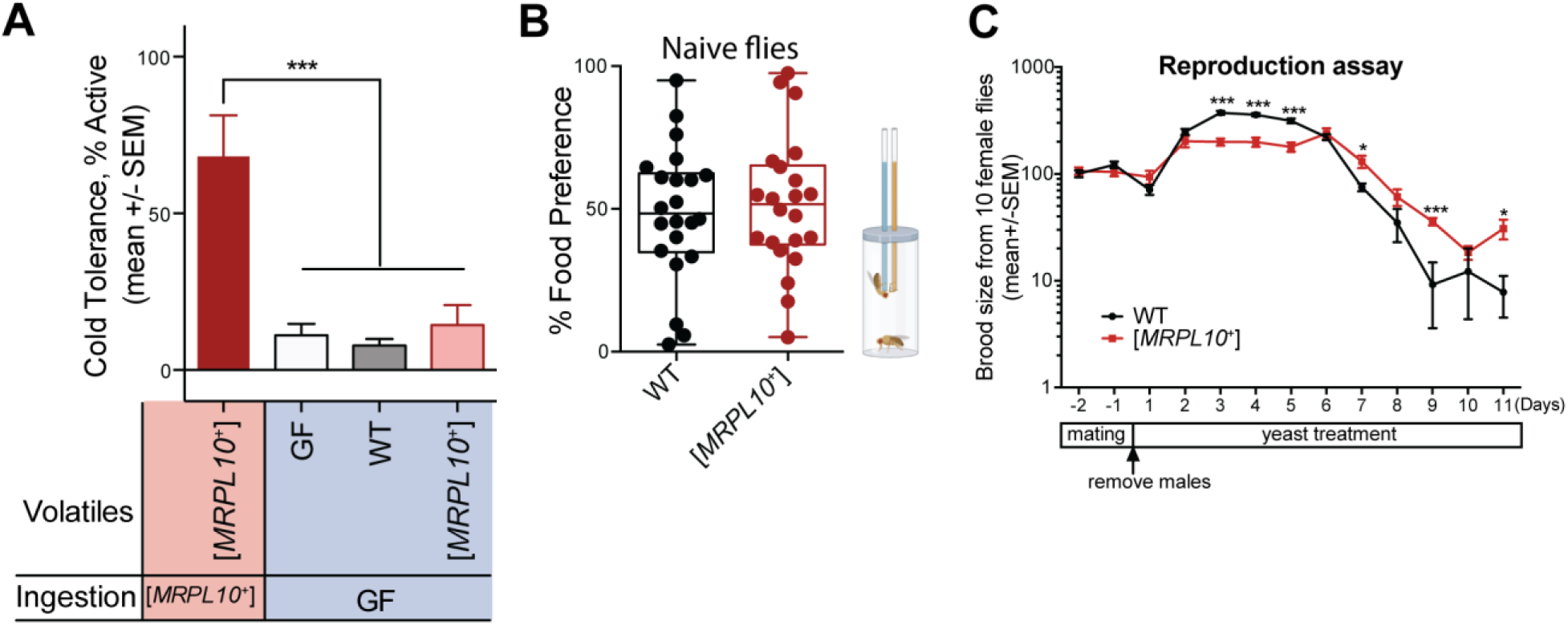
Volatile signaling, feeding behavior, and reproductive effects of [*MRPL10*^+^] yeast in *Drosophila melanogaster*. (A) Volatile cues emitted by [*MRPL10*^+^] yeast are not sufficient to induce the cold tolerance phenotype in flies. (B) Naïve flies (no prior exposure to live yeast) display no significant feeding preference between wild-type and [*MRPL10*^+^] yeast. (C) [*MRPL10*^+^] yeast reduce early brood size (days 3–5) but increased later brood size (days 7– 11) in flies, without a consistent overall trend. All data in (A)-(C) are based on three independent biological replicates.

**Supplementary Figure 3.**
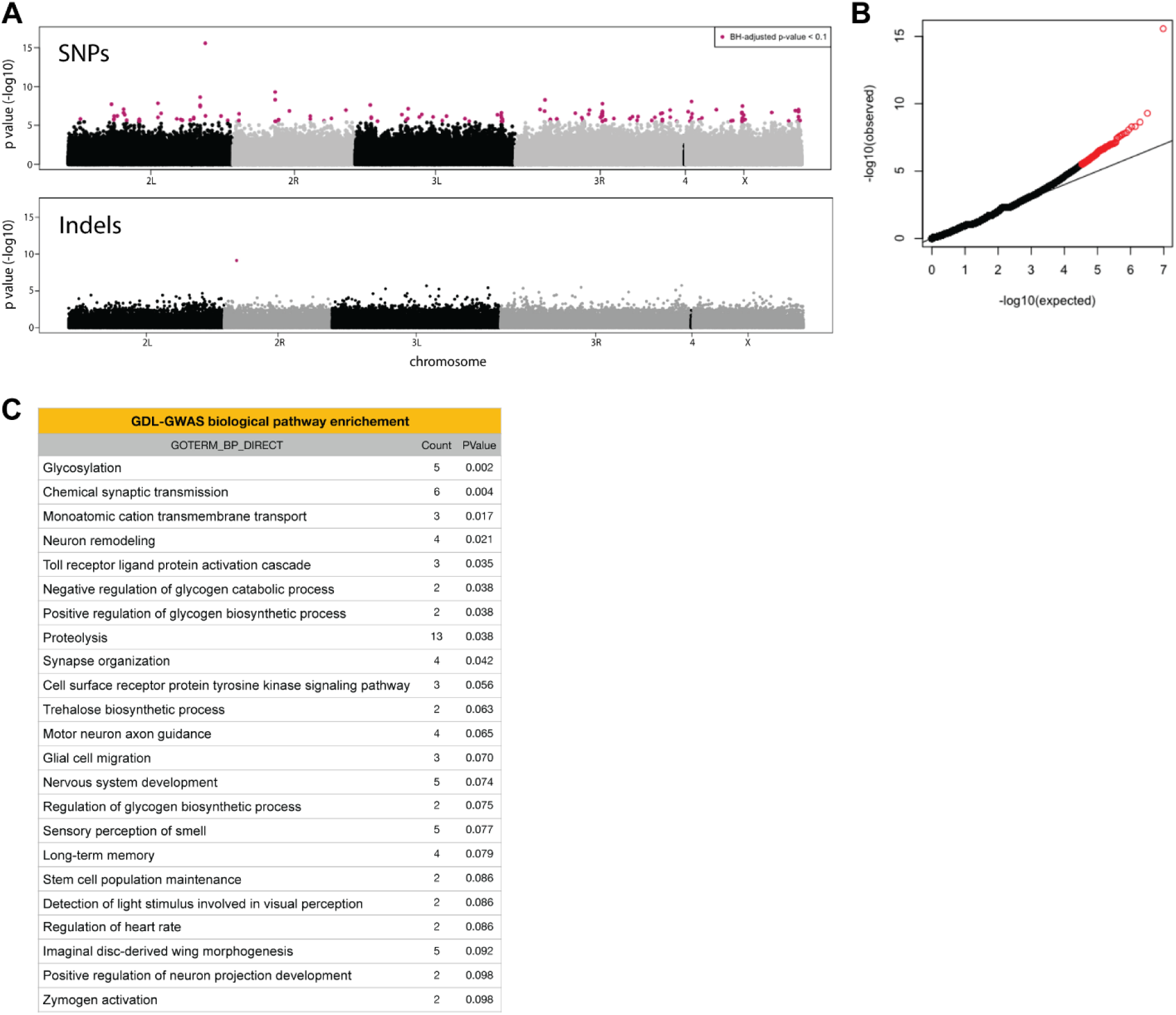
Genomic loci and pathway enrichment associated with [*MRPL10*^+^]-conferred cold tolerance in flies. (A) Manhattan plots depicting 144 single nucleotide polymorphisms (SNPs) and one insertion/deletion (indel) significantly associated with the cold tolerance phenotype conferred by [*MRPL10*^+^] yeast, based on genome-wide association analysis of 4,820,530 SNPs and 414,904 indels in GDL flies. Significant loci (Benjamini–Hochberg adjusted *p* < 0.1) are highlighted in red. (B) Quantile–quantile (Q–Q) plot comparing observed versus expected –log₁₀(*p*-values) from the genome-wide SNP association analysis. (C) Gene Ontology (GO) enrichment analysis of 246 candidate genes identified from GWAS reveals overrepresentation of metabolic and neuronal pathways.

**Supplementary Figure 4.**
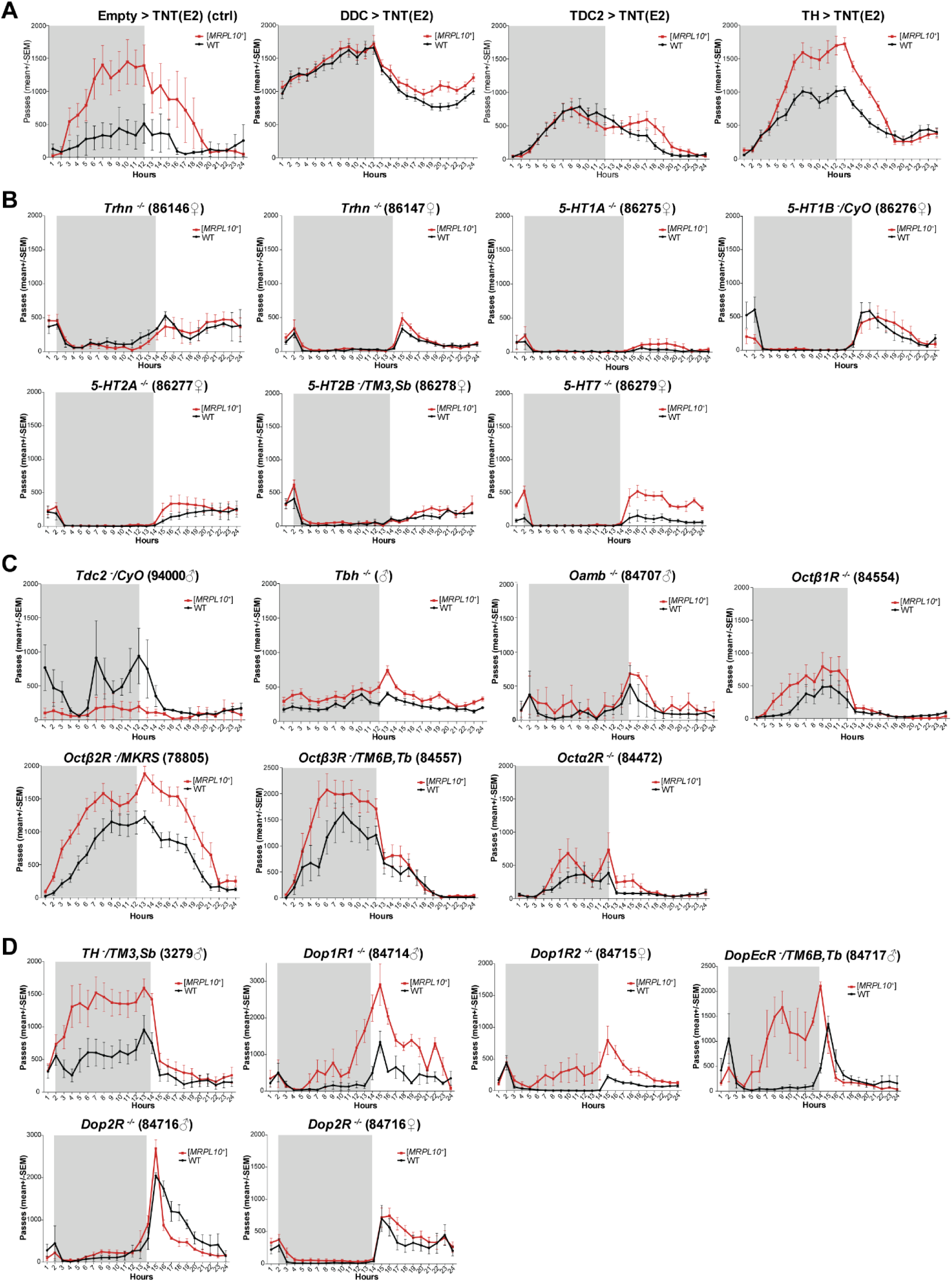
Serotonin and octopamine signaling mediate [MRPL10⁺]-induced locomotor activity. (A) Tissue-specific synaptic inhibition using tetanus toxin (neurotrapping) demonstrates that serotonergic and octopaminergic neurons are required for [*MRPL10*^+^]-induced increases in locomotion, while dopaminergic neurons are dispensable. (B) The serotonin biosynthesis gene *Trhn* and receptor genes *5-HT1A*, *5-HT1B*, *5-HT2A*, and *5-HT2B* are necessary for the locomotor response to [*MRPL10*^+^] yeast, whereas *5-HT7* is not. (C) Loss of the octopamine biosynthesis gene *Tdc2* or its receptors *Octβ2R* and *Octβ3R* triggers elevated locomotor activity even in flies fed wild-type yeast, mimicking the [*MRPL10*^+^] phenotype. (D) [*MRPL10*^+^]-induced locomotor activity is unaffected by loss of the dopamine biosynthesis gene *TH* or receptors *Dop1R1*, *Dop1R2*, and *DopEcR*, but is reduced in *Dop2R* mutants, suggesting a potential modulatory role for *Dop2R* in the full locomotor response. All data in (A)–(D) are based on at least three independent biological replicates.

Supplementary Table 1. Genetic variants associated with [*MRPL10*^+^]-induced cold tolerance identified through GWAS of *Drosophila* Global Diversity Lines (GDL). (UPLOADED AS A SEPARATE FILE)

**Supplementary Table 2.**
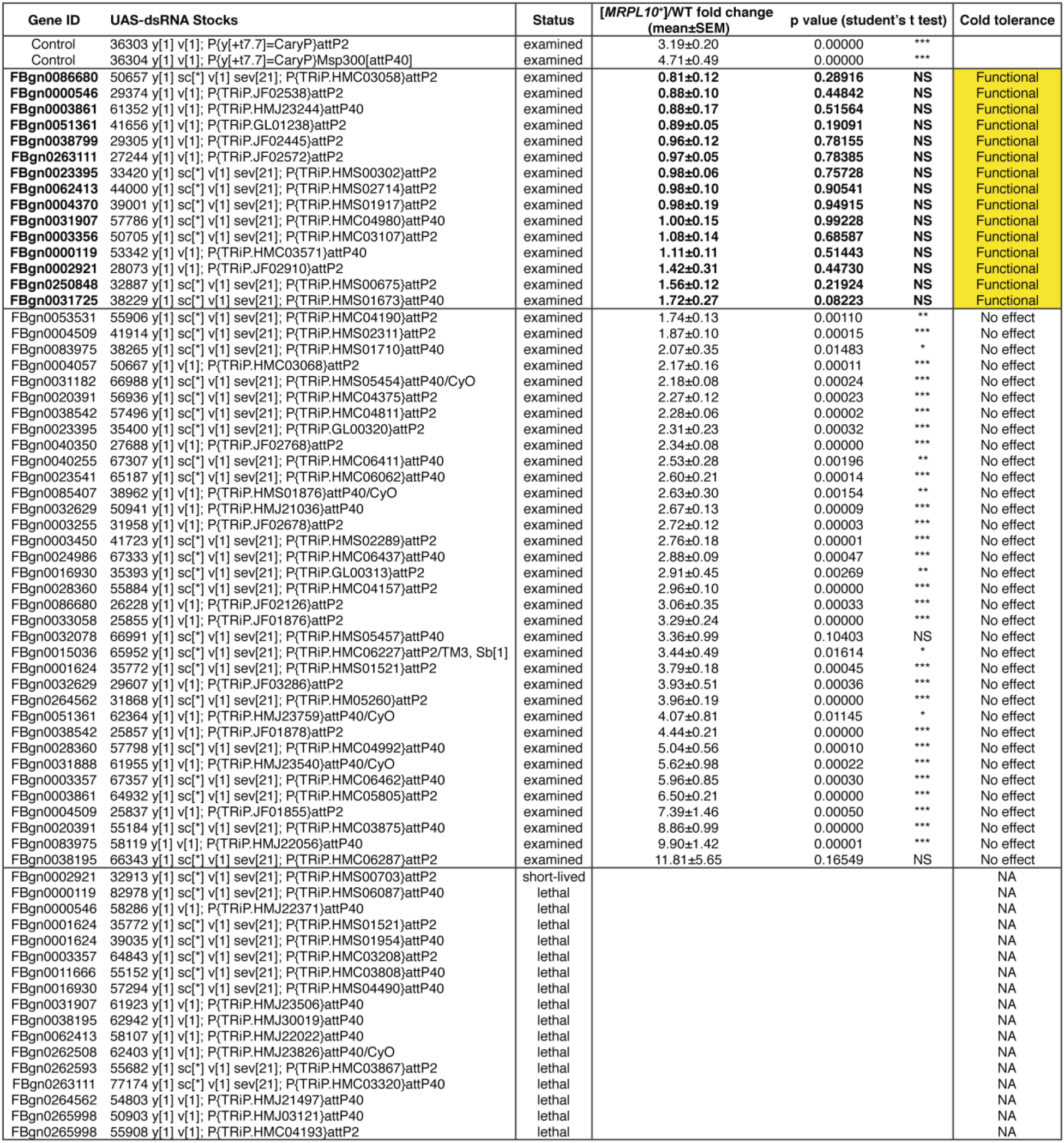
Genes and corresponding UAS-dsRNA lines (female) used in RNAi-based functional screening via crosses with GAL4 driver lines (male). A total of 15 genes were identified as required for the differential cold tolerance response to [*MRPL10*^+^] versus wild-type yeast.

## References

1. Alberti, S., Halfmann, R., King, O., Kapila, A., & Lindquist, S. (2009). A systematic survey identifies prions and illuminates sequence features of prionogenic proteins. Cell, 137(1), 146–158. 10.1016/j.cell.2009.02.044

2. Billeter, J. C., & Wolfner, M. F. (2018). Chemical Cues that Guide Female Reproduction in Drosophila melanogaster. J Chem Ecol, 44(9), 750–769. 10.1007/s10886-018-0947-z

3. Buser, C. C., Newcomb, R. D., Gaskett, A. C., & Goddard, M. R. (2014). Niche construction initiates the evolution of mutualistic interactions. Ecol Lett, 17(10), 1257–1264. 10.1111/ele.12331

4. Chakrabortee, S., Byers, J. S., Jones, S., Garcia, D. M., Bhullar, B., Chang, A., She, R., Lee, L., Fremin, B., Lindquist, S., & Jarosz, D. F. (2016). Intrinsically Disordered Proteins Drive Emergence and Inheritance of Biological Traits. Cell, 167(2), 369–381 e312. 10.1016/j.cell.2016.09.017

5. Chaudhary, V. B., Aguilar-Trigueros, C. A., Mansour, I., & Rillig, M. C. (2022). Fungal Dispersal Across Spatial Scales. Annual Review of Ecology Evolution and Systematics, 53, 69–85. 10.1146/annurev-ecolsys-012622-021604

6. Chen, Y. R., Ziv, I., Swaminathan, K., Elias, J. E., & Jarosz, D. F. (2021). Protein aggregation and the evolution of stress resistance in clinical yeast. Philos Trans R Soc Lond B Biol Sci, 376(1826), 20200127. 10.1098/rstb.2020.0127

7. Chernova, T. A., Wilkinson, K. D., & Chernoff, Y. O. (2014). Physiological and environmental control of yeast prions. FEMS Microbiol Rev, 38(2), 326–344. 10.1111/1574-6976.12053

8. Cho, H., & Rohlfs, M. (2023). Transmission of beneficial yeasts accompanies offspring production in Drosophila-An initial evolutionary stage of insect maternal care through manipulation of microbial load? Ecol Evol, 13(6), e10184. 10.1002/ece3.10184

9. Christiaens, J. F., Franco, L. M., Cools, T. L., De Meester, L., Michiels, J., Wenseleers, T., Hassan, B. A., Yaksi, E., & Verstrepen, K. J. (2014). The fungal aroma gene ATF1 promotes dispersal of yeast cells through insect vectors. Cell Rep, 9(2), 425–432. 10.1016/j.celrep.2014.09.009

10. Colinet, H., & Renault, D. (2014). Dietary live yeast alters metabolic profiles, protein biosynthesis and thermal stress tolerance of Drosophila melanogaster. Comp Biochem Physiol A Mol Integr Physiol, 170, 6–14. 10.1016/j.cbpa.2014.01.004

11. Du, Z., Park, K. W., Yu, H., Fan, Q., & Li, L. (2008). Newly identified prion linked to the chromatin-remodeling factor Swi1 in Saccharomyces cerevisiae. Nat Genet, 40(4), 460–465. 10.1038/ng.112

12. Gaspar, B. S., Rosu, O. A., Enache, R. M., Manciulea Profir, M., Pavelescu, L. A., & Cretoiu, S. M. (2025). Gut Mycobiome: Latest Findings and Current Knowledge Regarding Its Significance in Human Health and Disease. J Fungi (Basel), 11(5). 10.3390/jof11050333

13. Grangeteau, C., Yahou, F., Everaerts, C., Dupont, S., Farine, J. P., Beney, L., & Ferveur, J. F. (2018). Yeast quality in juvenile diet affects Drosophila melanogaster adult life traits. Sci Rep, 8(1), 13070. 10.1038/s41598-018-31561-9

14. Grenier, J. K., Arguello, J. R., Moreira, M. C., Gottipati, S., Mohammed, J., Hackett, S. R., Boughton, R., Greenberg, A. J., & Clark, A. G. (2015). Global diversity lines - a five-continent reference panel of sequenced Drosophila melanogaster strains. G3 (Bethesda), 5(4), 593–603. 10.1534/g3.114.015883

15. Halfmann, R., Alberti, S., & Lindquist, S. (2010). Prions, protein homeostasis, and phenotypic diversity. Trends Cell Biol, 20(3), 125–133. 10.1016/j.tcb.2009.12.003

16. Halfmann, R., Jarosz, D. F., Jones, S. K., Chang, A., Lancaster, A. K., & Lindquist, S. (2012). Prions are a common mechanism for phenotypic inheritance in wild yeasts. Nature, 482(7385), 363–368. 10.1038/nature10875

17. Hill, J. H., & Round, J. L. (2024). Intestinal fungal-host interactions in promoting and maintaining health. Cell Host Microbe, 32(10), 1668–1680. 10.1016/j.chom.2024.09.010

18. Huang, H., Wang, Q., Yang, Y., Zhong, W., He, F., & Li, J. (2024). The mycobiome as integral part of the gut microbiome: crucial role of symbiotic fungi in health and disease. Gut Microbes, 16(1), 2440111. 10.1080/19490976.2024.2440111

19. Jarosz, D. F., Lancaster, A. K., Brown, J. C. S., & Lindquist, S. (2014). An evolutionarily conserved prion-like element converts wild fungi from metabolic specialists to generalists. Cell, 158(5), 1072–1082. 10.1016/j.cell.2014.07.024

20. Jimenez-Padilla, Y., Adewusi, B., Lachance, M. A., & Sinclair, B. J. (2024). Live yeasts accelerate Drosophila melanogaster larval development. J Exp Biol, 227(19). 10.1242/jeb.247932

21. Jones, R., Fountain, M. T., Andreani, N. A., Gunther, C. S., & Goddard, M. R. (2022). The relative abundances of yeasts attractive to Drosophila suzukii differ between fruit types and are greatest on raspberries. Sci Rep, 12(1), 10382. 10.1038/s41598-022-14275-x

22. Korkmaz, Y., Belka, M., & Blumenstein, K. (2025). How cryptic animal vectors of fungi can influence forest health in a changing climate and how to anticipate them. Appl Microbiol Biotechnol, 109(1), 65. 10.1007/s00253-025-13450-0

23. Koyle, M. L., Veloz, M., Judd, A. M., Wong, A. C., Newell, P. D., Douglas, A. E., & Chaston, J. M. (2016). Rearing the Fruit Fly Drosophila melanogaster Under Axenic and Gnotobiotic Conditions. J Vis Exp(113). 10.3791/54219

24. Lancaster, A. K., Bardill, J. P., True, H. L., & Masel, J. (2010). The spontaneous appearance rate of the yeast prion [PSI+] and its implications for the evolution of the evolvability properties of the [PSI+] system. Genetics, 184(2), 393–400. 10.1534/genetics.109.110213

25. Lancaster, A. K., & Masel, J. (2009). The evolution of reversible switches in the presence of irreversible mimics. Evolution, 63(9), 2350–2362. 10.1111/j.1558-5646.2009.00729.x

26. Lazzaro, B. P., Flores, H. A., Lorigan, J. G., & Yourth, C. P. (2008). Genotype-by-environment interactions and adaptation to local temperature affect immunity and fecundity in Drosophila melanogaster. PLoS Pathog, 4(3), e1000025. 10.1371/journal.ppat.1000025

27. Libert, S., Zwiener, J., Chu, X., Vanvoorhies, W., Roman, G., & Pletcher, S. D. (2007). Regulation of Drosophila life span by olfaction and food-derived odors. Science, 315(5815), 1133–1137. 10.1126/science.1136610

28. Macmillan, H. A., & Sinclair, B. J. (2011). Mechanisms underlying insect chill-coma. J Insect Physiol, 57(1), 12–20. 10.1016/j.jinsphys.2010.10.004

29. Majumder, R., Sutcliffe, B., Taylor, P. W., & Chapman, T. A. (2020). Fruit host-dependent fungal communities in the microbiome of wild Queensland fruit fly larvae. Sci Rep, 10(1), 16550. 10.1038/s41598-020-73649-1

30. Mullinax, S. R., Darby, A. M., Gupta, A., Chan, P., Smith, B. R., & Unckless, R. L. (2025). A suite of selective pressures supports the maintenance of alleles of a Drosophila immune peptide. Elife, 12. 10.7554/eLife.90638

31. Murgier, J., Everaerts, C., Farine, J. P., & Ferveur, J. F. (2019). Live yeast in juvenile diet induces species-specific effects on Drosophila adult behaviour and fitness. Sci Rep, 9(1), 8873. 10.1038/s41598-019-45140-z

32. Ozcete, O. D., Banerjee, A., & Kaeser, P. S. (2024). Mechanisms of neuromodulatory volume transmission. Mol Psychiatry, 29(11), 3680–3693. 10.1038/s41380-024-02608-3

33. Patel, B. K., Gavin-Smyth, J., & Liebman, S. W. (2009). The yeast global transcriptional co-repressor protein Cyc8 can propagate as a prion. Nat Cell Biol, 11(3), 344–349. 10.1038/ncb1843

34. Quan, A. S., & Eisen, M. B. (2018). The ecology of the Drosophila-yeast mutualism in wineries. PLoS One, 13(5), e0196440. 10.1371/journal.pone.0196440

35. Rogoza, T., Goginashvili, A., Rodionova, S., Ivanov, M., Viktorovskaya, O., Rubel, A., Volkov, K., & Mironova, L. (2010). Non-Mendelian determinant [ISP+] in yeast is a nuclear-residing prion form of the global transcriptional regulator Sfp1. Proc Natl Acad Sci U S A, 107(23), 10573–10577. 10.1073/pnas.1005949107

36. Rosikon, K. D., Bone, M. C., & Lawal, H. O. (2023). Regulation and modulation of biogenic amine neurotransmission in Drosophila and Caenorhabditis elegans. Front Physiol, 14, 970405. 10.3389/fphys.2023.970405

37. Saad, S., & Jarosz, D. F. (2021). Protein self-assembly: A new frontier in cell signaling. Curr Opin Cell Biol, 69, 62–69. 10.1016/j.ceb.2020.12.013

38. Steck, K., Walker, S. J., Itskov, P. M., Baltazar, C., Moreira, J. M., & Ribeiro, C. (2018). Internal amino acid state modulates yeast taste neurons to support protein homeostasis in Drosophila. Elife, 7. 10.7554/eLife.31625

39. Suzuki, G., Shimazu, N., & Tanaka, M. (2012). A yeast prion, Mod5, promotes acquired drug resistance and cell survival under environmental stress. Science, 336(6079), 355–359. 10.1126/science.1219491

40. Tyedmers, J., Madariaga, M. L., & Lindquist, S. (2008). Prion switching in response to environmental stress. PLoS Biol, 6(11), e294. 10.1371/journal.pbio.0060294

41. Unckless, R. L., Howick, V. M., & Lazzaro, B. P. (2016). Convergent Balancing Selection on an Antimicrobial Peptide in Drosophila. Curr Biol, 26(2), 257–262. 10.1016/j.cub.2015.11.063

42. White, B. H., & Peabody, N. C. (2009). Neurotrapping: cellular screens to identify the neural substrates of behavior in Drosophila. Front Mol Neurosci, 2, 20. 10.3389/neuro.02.020.2009

43. Wickner, R. B. (1994). [URE3] as an altered URE2 protein: evidence for a prion analog in Saccharomyces cerevisiae. Science, 264(5158), 566–569. 10.1126/science.7909170

